# Stability-normalised walking speed: a new approach for human gait perturbation research

**DOI:** 10.1101/314757

**Authors:** Christopher McCrum, Paul Willems, Kiros Karamanidis, Kenneth Meijer

## Abstract

In gait stability research, neither self-selected walking speeds, nor the same prescribed walking speed for all participants, guarantee equivalent gait stability among participants. Furthermore, these options may differentially affect the response to different gait perturbations, which is problematic when comparing groups with different capacities. We present a method for decreasing inter-individual differences in gait stability by adjusting walking speed to equivalent margins of stability (MoS). Eighteen healthy adults walked on a split-belt treadmill for two-minute bouts at 0.4m/s up to 1.8m/s in 0.2m/s intervals. The stability-normalised walking speed (MoS=0.05m) was calculated using the mean MoS at touchdown of the final 10 steps of each speed. Participants then walked for three minutes at this speed and were subsequently exposed to a treadmill belt acceleration perturbation. A further 12 healthy adults were exposed to the same perturbation while walking at 1.3m/s: the average of the previous group. Large ranges in MoS were observed during the prescribed speeds (6-10cm across speeds) and walking speed significantly (P<0.001) affected MoS. The stability-normalised walking speeds resulted in MoS equal or very close to the desired 0.05m and reduced between-participant variability in MoS. The second group of participants walking at 1.3m/s had greater inter-individual variation in MoS during both unperturbed and perturbed walking compared to 12 sex, height and leg length-matched participants from the stability-normalised walking speed group. The current method decreases inter-individual differences in gait stability which may be beneficial for gait perturbation and stability research, in particular for studies on populations with different locomotor capacities.

## Introduction

Mechanical perturbations have been used for decades to investigate the stability of human walking (Berger et al., 1984; Marigold and Patla, 2002; Nashner, 1980; Quintern et al., 1985; Vilensky et al., 1999) and are now frequently investigated in falls prevention contexts (Gerards et al., 2017; Mansfield et al., 2015; Pai and Bhatt, 2007). In gait perturbation studies, self-selected walking speeds (e.g. Pai et al., 2014) or a prescribed walking speed for all participants (e.g. McCrum et al., 2016a) are commonly used, but each comes with drawbacks that complicate the interpretation of results. A prescribed walking speed will not result in comparable stability for all participants. This is problematic when comparing groups with different capacities during a gait perturbation task, as the relative challenge of the task will vary. Self-selected walking speeds, however, introduce other problems. Walking speed affects recovery strategy choice following slips (Bhatt et al., 2005) and trips (Krasovsky et al., 2014), the direction of balance loss following slipping (Smeesters et al., 2001) and differentially affects falls risk following tripping and slipping (Bhatt et al., 2005; Espy et al., 2010; Pavol et al., 1999). Gait stability at perturbation onset may also not be optimised at the self-selected speed and may differ across groups (Bhatt et al., 2005; Hak et al., 2013; Mademli and Arampatzis, 2014; Süptitz et al., 2012). The assessment of gait stability is also confounded by walking speed, which affects measures of dynamic gait stability using a centre of mass – base of support relationship model (Bhatt et al., 2005; Hak et al., 2013; Süptitz et al., 2012). Therefore, more sophistication in the choice of walking speed may be necessary for detailed study of reactive gait stability and adaptation processes.

Two possible solutions have been applied in previous gait perturbation studies. Two studies used 60% of the walk-to-run velocity to normalise the speed to participants’ neuromuscular capacities (Bierbaum et al., 2010, 2011) and a Froude number (a dimensionless parameter) for walking speed (Hof, 1996) has been applied to normalise the walking speed based on leg length (Aprigliano et al., 2016; Aprigliano et al., 2017; Martelli et al., 2013; Martelli et al., 2016). However, as these are normalisations based on a single parameter, neither of which are the sole determinants of gait stability, not all of the above described issues will be addressed. Therefore, further attempts to tackle these issues are warranted (McCrum et al., 2016b; McCrum et al., 2017).

Here, we present a new method for decreasing inter-individual differences in gait stability by normalising the walking speed based on gait stability. For this method we use the margins of stability (MoS) concept (Hof et al., 2005), one of the few well-defined and well-accepted biomechanical measures of mechanical stability of the body configuration during locomotion (Bruijn et al., 2013), useful for assessing changes in gait stability due to mechanical perturbations and balance loss. Additionally, we present results from a gait perturbation experiment comparing participants walking at their stability-normalised walking speed with participants walking all at the same prescribed speed.

## Methods

### Participants

Eighteen healthy adults participated in the first part of this study (eight males, 10 females; age: 24.4±2.5y; height: 174.9±7.4cm; weight: 74.6±15.2kg). Twelve healthy adults participated in the second part of the study (Table 1). The participants had no self-reported history of walking difficulties, dizziness or balance problems, and had no known neuromuscular condition or injury that could affect balance or walking. Informed consent was obtained and the study was conducted in accordance with the Declaration of Helsinki. The study protocol was approved by the Maastricht University Medical Centre medical ethics committee.

**Table 1:** Demographic characteristics of the participant groups in part two of the study.

|  | Sex | Age (y) | Height (cm) | Weight (kg) | Leg Length (cm) |
| --- | --- | --- | --- | --- | --- |
| 1.3m/s Group | 8 males, 4 females | 25.1±3.8 | 178.2±5.2 | 72.5±9.7 | 84.2±2.1 |
| Norm Group | 8 males, 4 females | 24.3±2.9 | 178.7±5.8 | 79±15.3 | 85.5±2.8 |
| Equivalent based on 90% CIs? | - | Yes | Yes | Yes | Yes |

### Setup and Procedures

The Computer Assisted Rehabilitation Environment Extended (CAREN; Motekforce Link, Amsterdam, The Netherlands), comprised of a dual-belt force plate-instrumented treadmill (Motekforce Link, Amsterdam, The Netherlands; 1000Hz), a 12-camera motion capture system (100Hz; Vicon Motion Systems, Oxford, UK) and a virtual environment that provided optic flow, was used for this study. A safety harness connected to an overhead frame was worn by the participants during all measurements. Five retroreflective markers were attached to anatomical landmarks (C7, left and right trochanter and left and right hallux) and were tracked by the motion capture system.

In the first part of the study (18 participants), the measurement sessions began with 60s familiarisation trials of walking at 0.4m/s up to 1.8m/s in 0.2m/s intervals. After approximately five to ten minutes rest, single two-to-three-minute-long measurements were then conducted at the same speeds. Following these measurements, the stability-normalised walking speed was calculated. To determine the stability-normalised walking speed, the mean anteroposterior MoS (see below) at foot touchdown of the final 10 steps of each walking trial (0.4m/s to 1.8m/s) were taken and fitted with a second order polynomial function. For each participant, the speed resulting in MoS of 0.05m was calculated. Based on our pilot testing, this value would result in walking speeds that would be possible for healthy adults of most ages. With certain populations, slower walking speeds would be required and then a greater MoS could be used. Participants then walked for three minutes at their stability-normalised walking speed, at the end of which, a gait perturbation was applied without warning. The perturbation consisted of a 3m/s2 acceleration of the right treadmill belt to 180% of the stability-normalised walking speed, thereby we also normalised the magnitude of the perturbation to the already normalised walking speed. The perturbation was triggered automatically by the D-Flow software of the CAREN, when the hallux marker of the to-be-perturbed limb became anterior to the stance limb hallux marker in the sagittal plane. The belt decelerated after toe-off of the perturbed limb.

In the second part of the study, 12 participants completed the same familiarisation protocol and then walked for three minutes at 1.3m/s (average stability-normalised walking speed of the 18 participants in the first part of the study). After this, they experienced the same treadmill belt acceleration perturbation. To compare these results with a matched sample, 12 participants from the first group of 18 were selected and matched specifically for sex, height and leg length to the participants in part two of the study (Table 1).

### Data Processing and Analysis

Marker tracks were filtered using a low pass second order Butterworth filter (zero-phase) with a 12Hz cut-off frequency. Foot touchdown was detected using a combination of force plate (50N threshold) and foot marker data (Zeni et al., 2008). The anteroposterior MoS were calculated at foot touchdown as the difference between the anterior boundary of the base of support (anteroposterior component of the hallux marker projection to the ground) and the extrapolated centre of mass as defined by Hof et al. (2005), adapted for our reduced kinematic model based on Süptitz et al. (2013). The MoS was calculated for: the final 10 steps of each set walking speed in the first part of the study; the mean MoS of the eleventh to second last step before each perturbation (Base); the final step before each perturbation (Pre); and the first recovery step following each perturbation (Post1).

### Statistics

A mixed effects model for repeated measures with walking speed as a fixed effect and Tukey post hoc comparisons was used to confirm a walking speed effect on the MoS. To determine whether a normalisation of walking speed based on body dimensions would assume equivalent gait stability, Pearson correlations between the stability-normalised walking speeds and participants’ height and leg length were conducted. A two-way repeated measures ANOVA with participant group (Stability-normalised walking speed [Norm] and 1.3m/s) and step (Base, Pre, Post1) as factors with post hoc Sidak’s tests for multiple comparisons were used to determine between group differences in the MoS. Equivalence tests using 90% confidence intervals were used to confirm the similarity of the groups’ demographics. Significance was set at α=0.05. Analyses were performed using Prism version 8 for Windows (GraphPad Software Inc., La Jolla, California, USA).

## Results and Discussion

Walking speed significantly affected the MoS (F[2.547, 42.93]=1485, P<0.0001; Fig. 1) and Tukey’s multiple comparisons tests revealed significant differences for each speed compared to all other speeds (P<0.0001; Fig. 1). These results agree with previous work (Bhatt et al., 2005; Hak et al., 2013; Süptitz et al., 2012). A range of MoS values were observed for each speed (approximately 6-10cm), even among these healthy participants, confirming some of the issues related to prescribed walking speeds in gait stability research discussed above. The strong relationship between walking speed and MoS also has relevance for clinical studies conducting self-paced gait measurements with an assessment of gait stability. Patients who improve in walking speed may demonstrate a reduction in MoS, which may not be reductions in the stability of the patients’ gait *per se*, but simply an artefact of the improved walking speed.

**Fig. 1:**
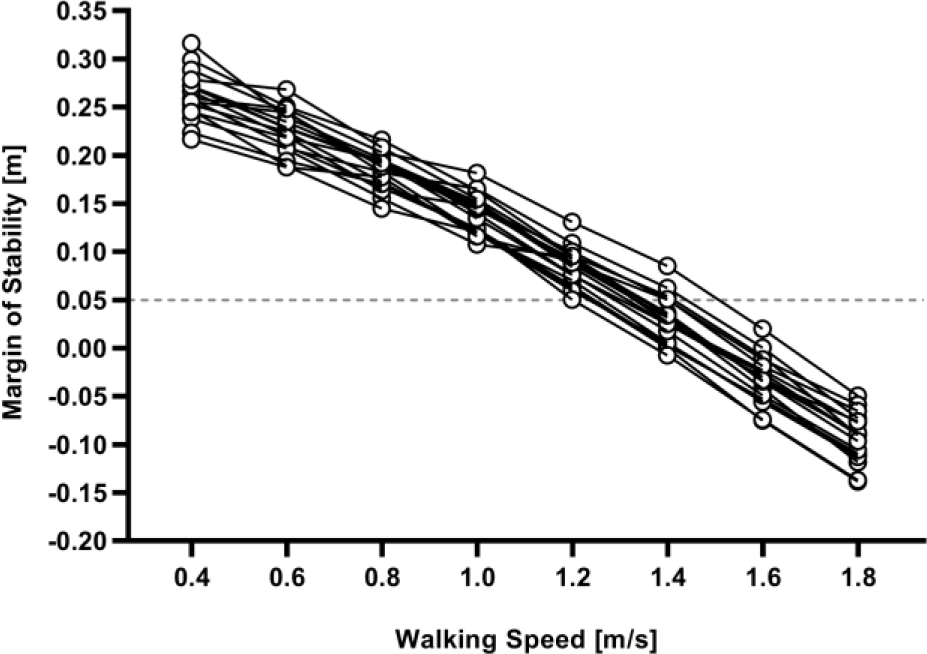
Individual margins of stability at foot touchdown over the different walking speeds. The dashed line represents the margin of stability used to determine the stability-normalised walking speed.

The stability-normalised walking speeds (range from 1.22m/s to 1.51m/s with a mean±SD of 1.3±0.1m/s) resulted in MoS very close to the desired outcome of 0.05m (within one SD of the mean MoS for 15 of the 18 participants; Fig. 2A). The stability-normalised walking speed also reduced between-participant variability in MoS (as shown by the group level standard deviations; Fig. 2B). These combined results indicate that the stability-normalisation was successful in reducing between participant differences in MoS during walking, even in a homogenous group of healthy young adults.

**Fig. 2:**
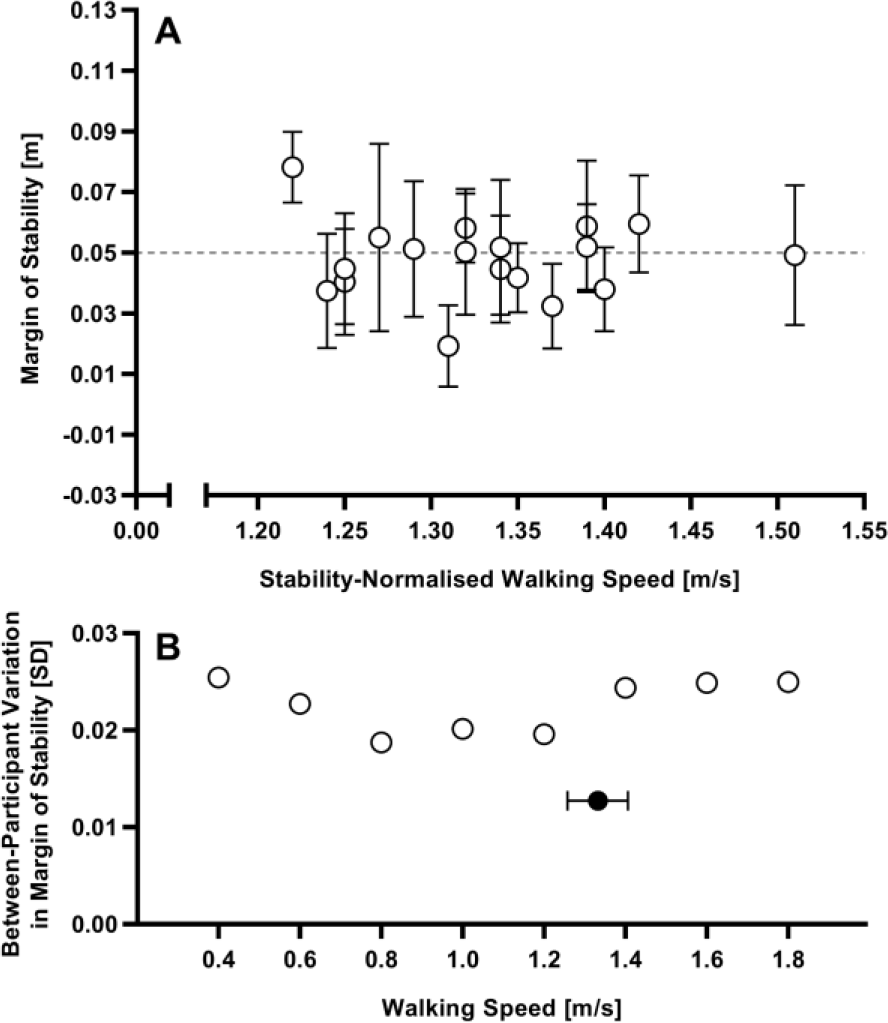
**A:** Means and standard deviations of the margins of stability at touchdown of the final 10 steps at the stability-normalised walking speed for each individual participant. The desired MoS of 0.05m at foot touchdown is indicated by the dashed line. **B:** The between-participant variation in the margins of stability (standard deviation at group level) for the final 10 steps at each walking speed (the stability-normalised walking speed trials are indicated with the black circle; mean and standard deviation).

Small, non-significant correlations between the determined stability-normalised walking speeds and the participants’ height and leg length were found (Fig. 3). The outcomes of our correlation analysis suggest that height and leg length did not significantly affect the calculation of stability-normalised walking speed, suggesting that a normalisation of walking speed based on body dimensions does not assume equivalent gait stability, at least not when assessed by the MoS concept.

**Fig. 3:**
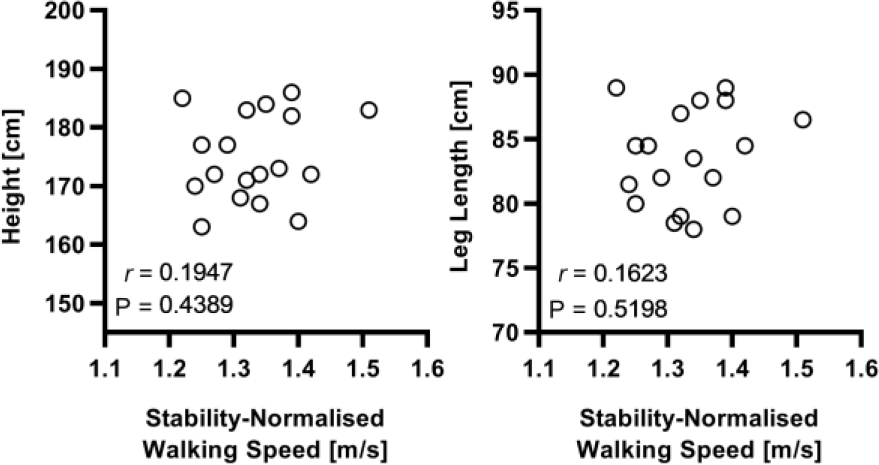
Pearson correlations between the participants’ stability-normalised walking speeds and their height and leg length.

For the second part of the study, the 12 participants were successfully matched to the 12 of the 18 participants from part one of the study (Table 1). During the perturbations, the 1.3m/s group had a greater range in MoS values during Base, Pre and Post1 (Fig. 4). A two-way repeated measures ANOVA revealed a significant effect of group (F[1, 22]=6.409, P=0.019), step (F[1.097, 24.14]=8.34, P=0.0068) and a significant group by step interaction (F[2, 44]=15.4, P<0.0001) on MoS. Sidak post hoc tests revealed a significant difference between Norm and 1.3m/s groups at Post1 (P=0.0049). While part of the differences found may be due to chance, the current comparison suggests that the stability-normalised walking speed and the normalised perturbation (acceleration to a peak speed 180% of the walking speed) reduce the inter-individual differences in MoS during both unperturbed and perturbed walking, at least with the current protocol. The significant difference found at Post1 between the groups also aligns with the previous studies reporting different responses to perturbations experienced while walking at different speeds (Bhatt et al., 2005; Krasovsky et al., 2014).

**Fig. 4:**
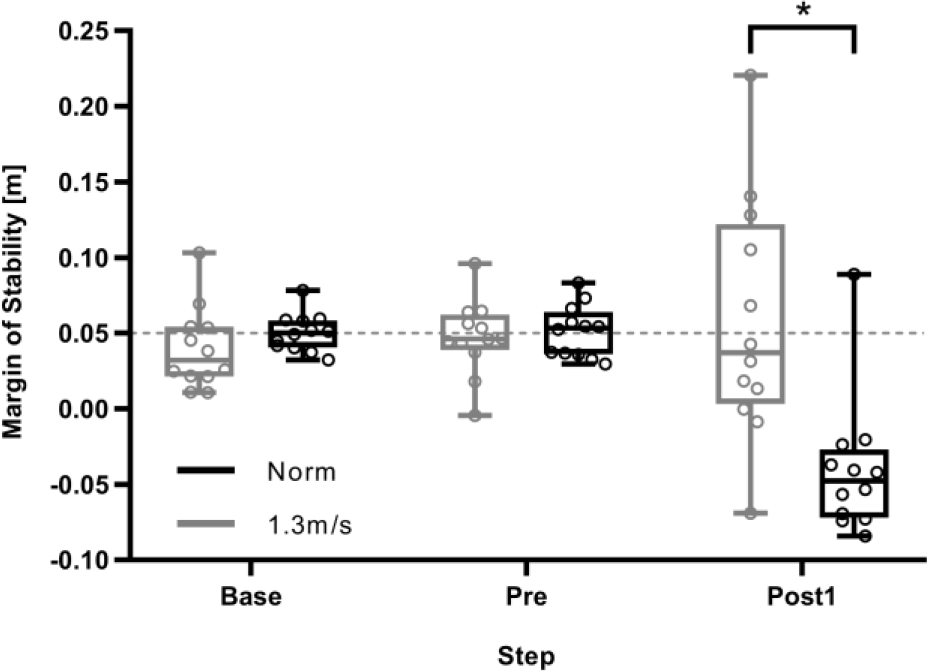
Margins of stability during unperturbed and perturbed walking of participants walking at their stability-normalised walking speed (Norm) and participants walking at 1.3m/s. Base: the mean MoS of the eleventh to second last step before each perturbation; Pre: the final step before each perturbation; Post1: the first recovery step following perturbation. *: Significant difference (Sidak post hoc test: P=0.0049).

As the MoS – walking speed relationship from 1.0-1.6m/s appeared to be linear in part one of the study (Fig. 1), a simple linear regression was calculated for 1.0-1.6m/s. A significant regression equation was found (Fig. 5). Future research could use this (or similar) as a simple, efficient method for increasing the dynamic similarity in gait stability across participants, by measuring participants walking at a single speed from 1.0-1.6m/s and using this equation to prescribe speeds that would result in similar MoS values. It is, however, worth highlighting that the current participants were young healthy adults; the walking speed – MoS relationship may be altered in other populations.

**Fig. 5:**
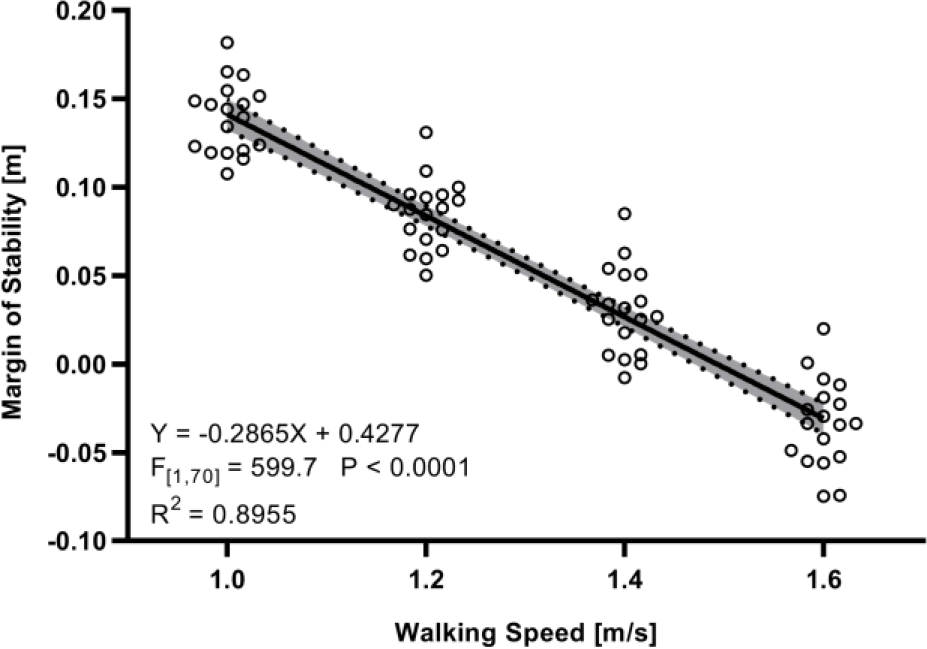
Margins of stability as a function of walking speed between 1.0 and 1.6m/s. The shaded area represents the 95% confidence intervals of the regression line.

In conclusion, large ranges in MoS were observed and walking speed significantly affected MoS even within these young healthy participants, confirming some issues related to walking speed choice in gait stability research. The current methods reduced between-participant variability in MoS during both unperturbed and perturbed walking, meaning that the method could be beneficial for gait stability studies comparing groups with different locomotor capacities. An equation has been provided that can be used following a single gait trial to increase the dynamic similarity of gait stability between participants.

## Acknowledgments

CM was funded by the Kootstra Talent Fellowship awarded by the Centre for Research Innovation, Support and Policy (CRISP) and by the NUTRIM Graduate Programme, both of Maastricht University Medical Center+. The authors thank Julia Agmon for assistance with the measurements.

